# Graph-pMHC: Graph Neural Network Approach to MHC Class II Peptide Presentation and Antibody Immunogenicity

**DOI:** 10.1101/2023.01.19.524779

**Authors:** William John Thrift, Jason Perera, Sivan Cohen, Nicolas W. Lounsbury, Hem Gurung, Chris Rose, Jieming Chen, Suchit Jhunjhunwala, Kai Liu

## Abstract

Antigen presentation on MHC Class II (pMHCII presentation) plays an essential role in the adaptive immune response to extracellular pathogens and cancerous cells. But it can also reduce the efficacy of large-molecule drugs by triggering an anti-drug response. Significant progress has been made in pMHCII presentation modeling due to the collection of large-scale pMHC mass spectrometry datasets (ligandomes) and advances in deep machine learning. Here, we develop graph-pMHC, a graph neural network approach to predict pMHCII presentation. We derive adjacency matrices for pMHCII using Alphafold2-multimer, and address the peptide-MHC binding groove alignment problem with a simple graph enumeration strategy. We demonstrate that graph-pMHC dramatically outperforms methods with suboptimal inductive biases, such as the multilayer-perceptron-based NetMHCIIan-4.0 (+22.84% average precision). Finally, we create an antibody drug immunogenicity dataset from clinical trial data, and develop a method for measuring anti-antibody immunogenicity risk using pMHCII presentation models. In comparison with BioPhi’s Sapiens score, a deep learning based measure of the humanness of an antibody drug, our strategy achieves a 7.14% ROC AUC improvement in predicting antibody drug immunogenicity.

## Introduction

Major histocompatibility complex class II (MHCII) molecules play an essential role in the immune system’s defense against pathogens. Peptides presented on MHCII (pMHCII) are primarily derived from extracellular proteins, and pMHCII can then be recognized by CD4+ helper T lymphocytes (CD4 T cells) to stimulate cellular and humoral immunity.^1^ Due to the complexity of determining which peptides are likely to be presented by MHCII, deep learning based computational tools have been developed and used extensively for various applications. _2–5_

Three main strategies have emerged for modeling: 1) multilayer perceptrons (MLP), exemplified by NetMHCIIpan-4.0,^6^ 2) sequence based models, such as the transformer based MHCAttnNet,^7^ and 3) convolutional neural networks (CNN), such as PUFFIN.^8^ Each of these approaches: MLP, RNN, and CNN, bring their own inductive bias^9^ – assumptions made in the design of the model – to the pMHCII prediction task. Graph neural networks (GNN) better capture real protein systems by leveraging prior knowledge of edges that connect and show interactions between nodes (residues). Indeed, GNNs were recently applied to pMHCII presentation, but faced limited success, only approaching the performance^10^ of NetMHCIIpan-4.0 in class II. These results show the risk of over-reliance on protein folding models, especially when we can capitalize on the vast amount of peptidome data available for training, and the unique features of the pMHCII interaction that can be leveraged for better performance.

We are particularly interested in the application of these models to biotherapeutic (particularly antibody drugs) deimmunization. Anti-drug antibodies (ADA), which the immune system uses to clear biotherapeutics, reduce the safety of biotherapeutics by 60%, and similarly 40% of biotherapeutics report a reduction in efficacy.^11^ Traditionally, humanization strategies like complementarity-determining region (CDR) grafting are used to reduce ADAs^12^ but require expert knowledge. Deep learning-based humanization has recently been shown to be predictive of antibody ADA responses, but primarily for mouse-derived antibody drugs.^13^ pMHCII prediction models too have been used to deimmunize proteins,^14–16^ but a high quality dataset of similar biotherapeutics is lacking to assess these disparate deimmunization methods.

In this work, we introduce graph-pMHC, a pMHCII peptide presentation model with state-of-the-art performance. We use Alphafold2-multimer (AF2)^17^ to generate canonical pMHC adjacency matrices for the MHCII binding grooves, and an alignment-based approach to learn the correct binding core location within a peptide by enumerating possible graphs. We demonstrate that this strategy dramatically outperforms current literature methods (21.47% average precision), and has a near optimal inductive bias. To assist this analysis, we develop a new test-train split strategy that ensures even distributions of ontologies and k-mers. Finally, we introduce a new biotherapeutic immunogenicity dataset with the ADA rate of 107 antibodies, which enables a comparison of pMHCII-based immunogenicity risk assessment with that of humanness-based methods, with our strategy outperforming SAPIENS^13^ by 7.41% ROC AUC. This work represents both a large step forward in the performance of pMHCII presentation models, as well as introducing new, high quality datasets for them to be compared in the future.

## Results

### Alphafold2-predicted pMHC residue interactions a concordant with crystal structures

We first identify the adjacency matrices for pMHCII using alphafold2-multimer^17^ (AF2) to predict the graph structure of pMHCII residue interactions. This is necessary due to the limited peptide and allele diversity (36 unique peptides, and 32 unique alleles) in the available empirical crystal structures, which is dramatically less than the 1000s of allotypes registered on the Immunogenetics information System (IMGT), and the 10,000s of unique peptides measured in pMHCII ligandomes. Indeed, no crystal structures exist for DQ with a non-covalently linked peptide. As AF2 is too computationally expensive at the time of writing to obtain predictions for all pMHCII in our dataset, we sought to obtain one canonical adjacency matrix for each gene, DR, DP, and DQ. Thus we must determine if adjacencies are conserved across alleles, peptide sequences, and peptide lengths. Figure 1 b) depicts the derived adjacency matrices corresponding to the empirical crystal structure and the AF2-solved structure in Figure 1 a). One may observe good correspondence between the AF2 structure and the empirical structure, with somewhat more adjacencies predicted by AF2, but only 1 missed adjacency. Below we discuss aggregate results on available PDB structures.

**Figure 1.**
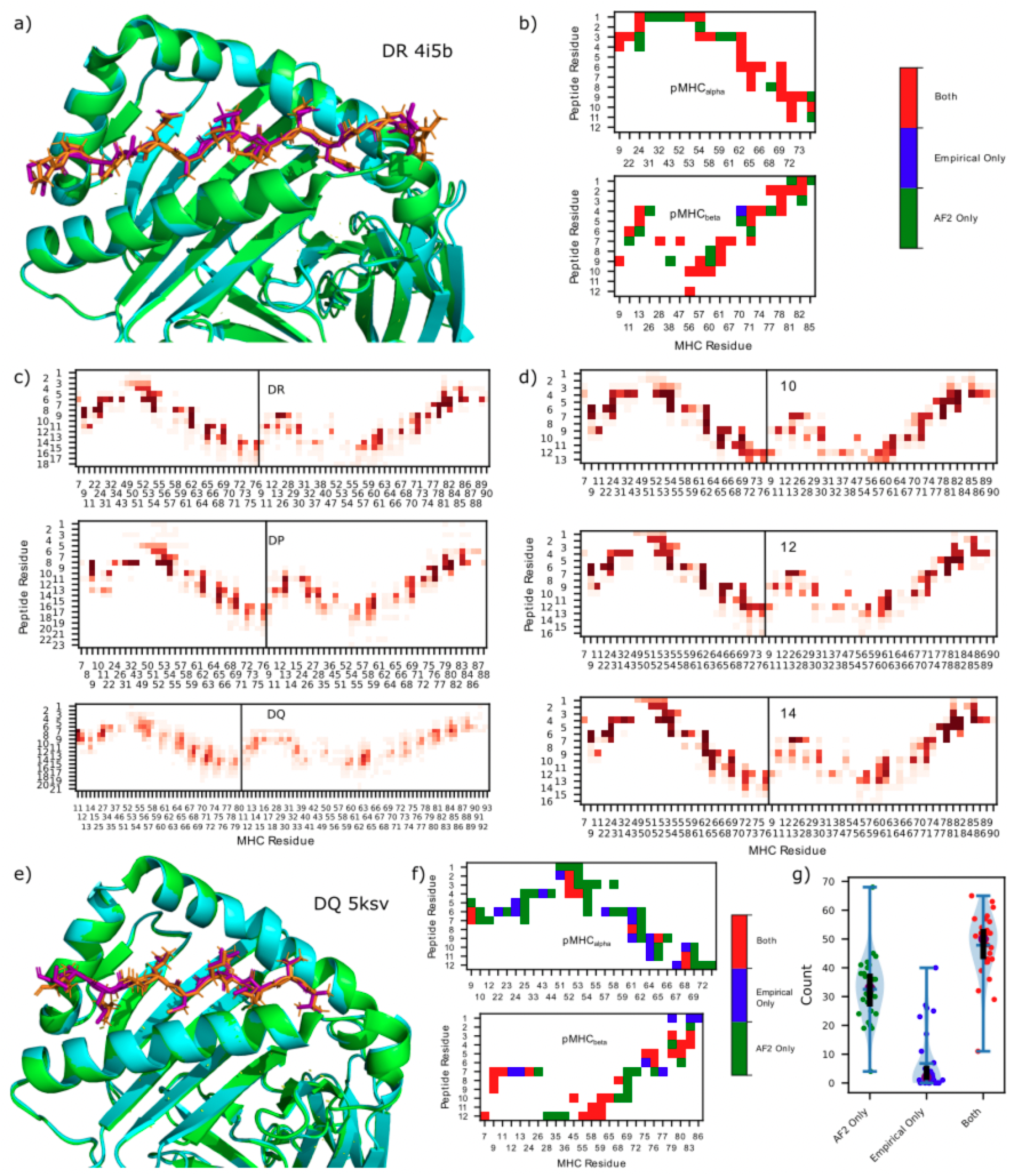
AF2 derived adjacency matrix. a) 3d structure of empirical crystal structure (PDB ID: 4i5b) of pMHCII (cyan, purple) and AF2 predictions (green, orange) for 4i5b, HLA-DRA*01:01/HLA-DRB1* VVKQNCLKLATK. b) Contact map (<4 Å) between pMHCII for the empirical crystal structure and the AF2 predictions corresponding to the pMHC in a). c) DR (upper) (166 adjacencies), DP (57 adjacencies) (center), and DQ (89 adjacencies) (lower) contact map aggregates over various lengths and alleles. d) Contact map for 10mer (upper) (18 adjacencies), 12mer (center) (17 adjacencies), and 14mer (lower) (22 adjacencies) peptides predicted by AF2 for various alleles. e) 3d structure of empirical crystal structure of pMHCII (cyan, purple) and AF2 predictions (green, orange) for 5ksv, HLA-DQA1*05/DQB1*02 MATPLLMQALPMGAL. f) contact map (<4 Å) between pMHCII for empirical crystal structure and AF2 predictions corresponding to the pMHCII in e). g) Violin plot of the adjacencies found only by AF2, only in the empirical source, and both, plotted with green, blue, and red dots, respectively for each PDB structure. The blue horizontal bar represents the median value, the purple dot represents the median value, the black line runs from the 1st to 3rd quartile.

Using this approach we obtained AF2 structure predictions for three different peptides (see methods for peptide selection criteria) for each allele in our presentation dataset (discussed below), and plot the adjacencies in Figure 1 c). As only a subset of peptide residues dock into the binding groove of MHC (the binding core, usually assumed to be 9 residues),^18^ we align the peptides relative to MHC (see methods). For DR and DP genes (DQ discussed below), we observe many highly conserved (yet distinct for DR/DP) adjacencies, and thus use them for the canonical adjacency matrices. Interestingly, we observe (Supplemental Figure 1) a median of 13 peptide residues in contact with MHC.

For the one adjacency per gene approach to be valid, the adjacency of peptides with different lengths as well as different alleles must be conserved across different structures. This is particularly interesting for peptides with fewer residues than are typically in contact with MHC. In Figure 1 d) we show aggregate adjacencies across 33 DR alleles for lengths 10, 12, and 14. We find that even in this case, the conserved adjacencies found before are again conserved for these short peptides.

From Figure 1 c) we observe significantly more variation in the adjacencies (as well as adjacency differences) for DQ alleles than DR and DP. Yet, comparison of an AF2 structure with the empirical crystal structure shown in Figure 1 e) shows good agreement. The contact map in Figure 1 f) again shows overprediction of AF2 contacts, but now with somewhat more adjacencies missed by AF2. DQ is an interesting case in pMHCII presentation, with models generally performing worse than DR and DP, the large variations in contact may be a contributing factor to this behavior.

Analyzing 43 pMHCII structures obtained from the PDB, we observe general agreement, with the median number of adjacencies found exclusively in the empirical sources to be 2. Figure 1 g) depicts violin plots of the (dis)agreement between empirical sources and AF2. Surprisingly, AF2 routinely places peptides binding in the opposite orientation (eg C-N instead of N-C), with 11 of the 43 structures placed in this orientation. While some studies suggest that these are possible orientations,^19,20^ none were in agreement with pdb structures, and so all the opposite-orientation peptides are filtered out prior to our creation of a canonical adjacency matrix (and are not plotted in Figure 1 g)). We also find AF2 placing 1 of the 43 structures in a non-flat orientation, we similarly filter out all non-flat peptides for determining the canonical adjacency matrix. Finally, AF2 predicts 1 peptide’s binding core starting position incorrectly, yielding bad predictions for adjacency, as observed as an outlier in Figure 1g).

### The Graph-MHC model

Figure 2 a) depicts a schematic of the graph structure of the pMHCII model used in this work. Two definitions of the peptide binding core were evaluated, the 9mer binding core defined by the anchor residues, and the full 15mer binding core identified using crystal structures (see figure 4 b)). Both the peptide binding core, as well as the peptide flanking residues (which are not in contact with MHCII) are included into the graph. Similar to other work,^6,18^ we use a pseudosequence to represent the MHC, where only MHC residues adjacent to the peptide are included. Importantly, we take advantage of edge features to inform the model of the type of interactions between residues, namely intermolecular interactions (those between peptide and MHC), intramolecular interactions (those within a peptide, MHC, or flank sequence), and division of protein flanks (between peptide and flank sequences). By connecting these sequences with its own unique edge token, we are able to create an end-to-end model of the entire presentation pathway.

**Figure 2.**
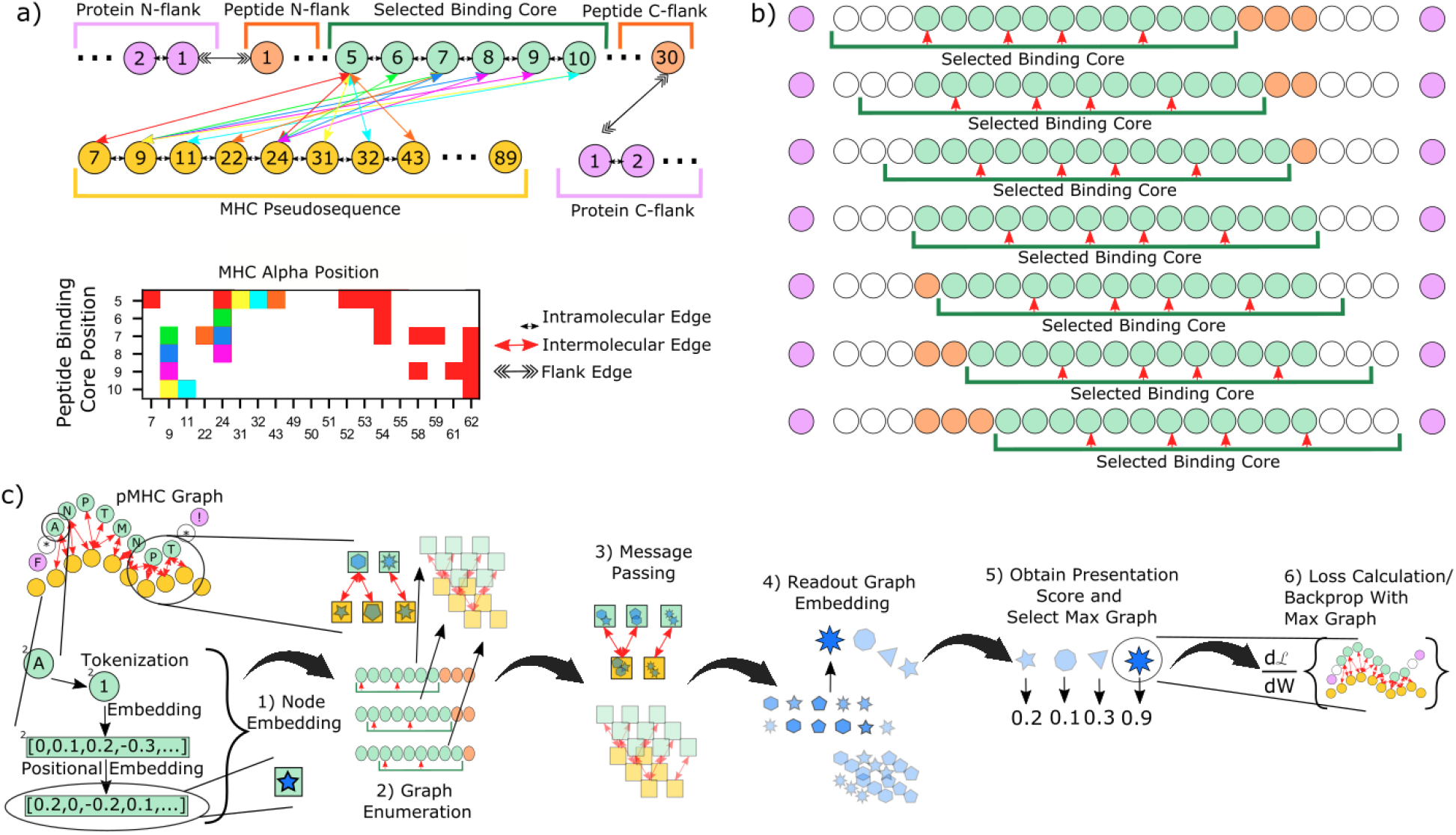
graph-pMHC scheme. a) Schematic of the graph generation approach used in this work. Intermolecular edges in the graph (upper), are color-matched with the adjacency matrix (lower) for clarity. Different edge types are used for intermolecular, intramolecular, and flank edges. Circles represent graph nodes, which are amino acid residues from the peptide, MHC, and flank sequences. b) Schematic of the ‘graph enumeration’ procedure used in this work. A green bar below the peptide represents the 15mer binding core, while red arrows denote the anchor residues. The 9mer binding core used in this work runs from the first anchor to the final anchor. Green circles represent the selected binding core, orange circles represent the residues in the peptide flanking the binding core, and purple circles represent flanking residues obtained from the protein from which the peptide was derived. White circles represent dummy residues used to enable 15mer binding core selection to begin or end at the first or last anchor residue. c) Schematic of graph-pMHC, the GNN model framework used. Circles represent amino acid residues, squares and various shapes represent vector node embeddings.

In general, a peptide may be larger than the size of the binding core, thus requiring an alignment procedure to select the residue where the binding core starts. Here we enumerate all binding core starting positions with a sliding window, which we refer to as ‘graph enumeration’, and depict in Figure 2 b). As anchors drive the binding to MHC, we allow for the 15mer binding core to begin and end at anchor residue, while for smaller binding cores, dummy residues are used for padding to obtain 15mers. Most pMHCII presentation data comes from mutli-allotypic (MA) samples, and so we perform an additional enumeration process across all possible peptide-allotype pairs.

Our general model framework, graph-pMHC, is depicted in Figure 2 c). First, residues are tokenized and embedded with a learned lookup table. Then, we apply a positional encoding using a learned lookup table for each position. The use of positional encodings for GNNs is uniquely applicable to our domain, adding extra information about the absolute position of the binding core. Next, graph enumeration is performed, generating all possible peptide-allotype and binding core starting position graphs (Supplemental Figure 2 depicts the convergence behavior of these selections). Typical GNN message passing is then performed separately for each graph to update the residue embeddings. Subsequently, a graph readout strategy is used to get a single vector embedding for the graph. In the ablation study below, we show how the choice of message passing and graph readout strategies impact performance. Finally, fully-connected layers map these graph embeddings to a presentation likelihood, such that the allotype-peptide-binding core with the best likelihood is taken as the canonical graph for the sample, and is used to calculate loss and perform backpropagation.

### Train/test split strategy to minimize gene ontology biases

In order to train our model, we have assembled large pMHCII ligandomes that have recently been published across 9 studies (Figure 3 a). In total, we aggregate 527,302 peptide:genotype pairs, with 250,643 unique peptides, and 75 unique alpha and beta MHCII chains. The data is heavily skewed towards MA, with 408,111 MA peptide:genotype pairs, and 119,191 single allotypic peptide:genotype pairs.

**Figure 3.**
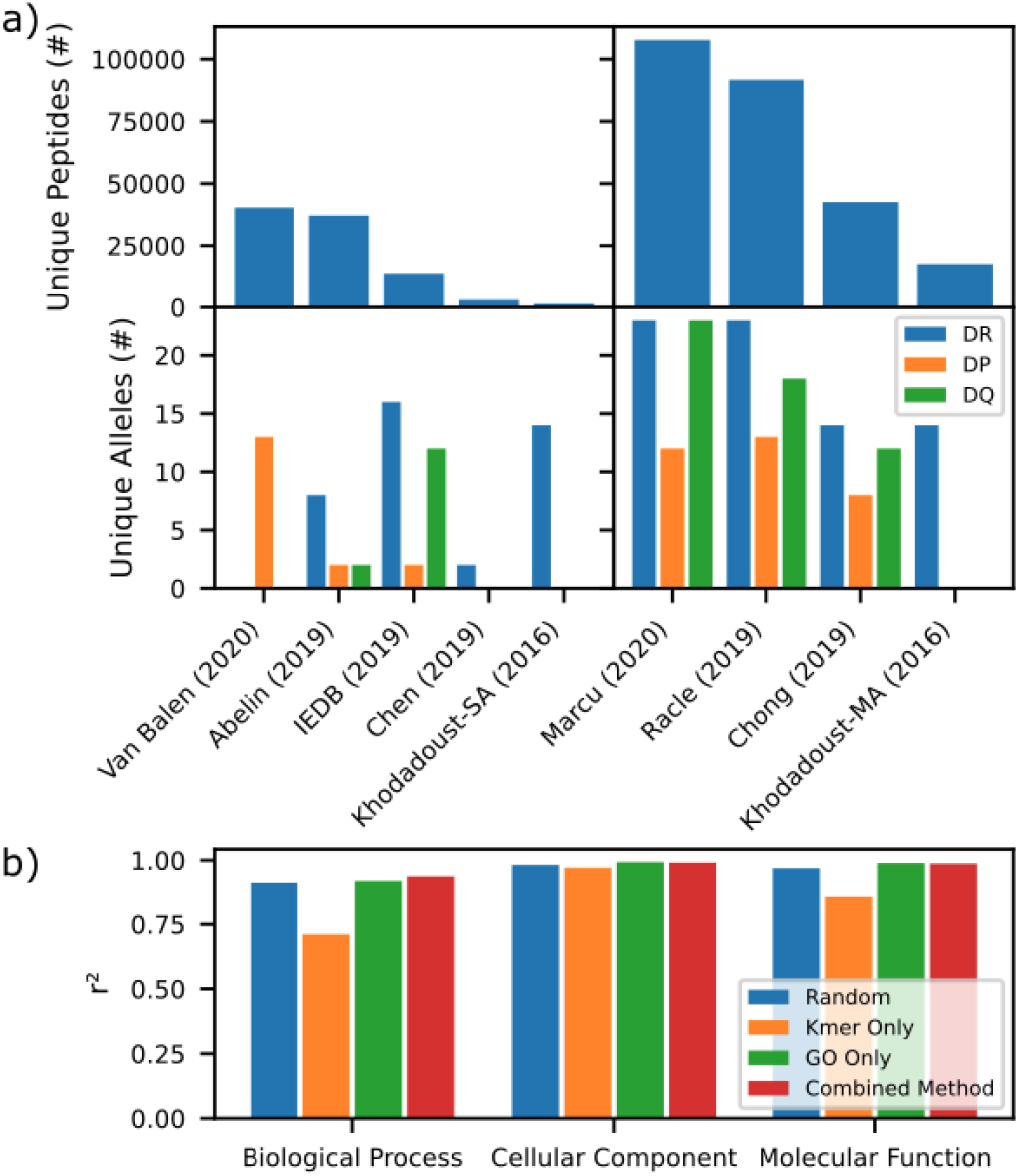
pMHCII dataset. a) Barchart depicting the number of unique peptides (upper) and unique alleles (lower) for monoallelic (left) and multiallelic sources (right). b) Barchart depicting goodness of split of genes with various biological processes, cellular components, and molecular function between our train and test datasets, as measured by r^2^ for various test-train splitting methods (random, kmer, gene ontology, and our combined method).

We desire a test/train split method that best reflects the model’s true performance. Others have already noted the importance of ensuring that 9mer overlap between test and train will skew test performance higher.^18^ Yet, little focus has been placed on ensuring good splitting of various gene ontologies into test and train, which could lead to misleading performance metrics for underrepresented ontologies in the test data. To this end, we developed a method combining 9-mer overlap and gene ontology awareness to reduce 9-mer overlap between train and test while maintaining a relatively even distribution of genes with various cellular localizations (cc), molecular functions (mf), and biological processes (bp) across train and test (see methods). Figure 3 b) depicts the quality of split (measured by r^2^) for the various splitting methods. One may observe that the traditional kmer overlap strategies for producing test-train splits leads to different ccs, mfs, and bps being over represented in either test or train compared to a random selection. A gene ontology (GO) based split solves the issue, but leaves identical 9mers in both train and test. Our combined strategy achieves much of the GO splitting while only allowing minimal 9mer overlap, with 0.35% overlap compared to 0.0029% overlap for the kmer-only strategy and 2.09% overlap compared to a random split.

### Investigating the Optimal Model Architecture and Adjacency Matrix

Figure 4 a) depicts performance (measured using average precision (AP)) of baseline graph-pMHC on our pMHCII presentation dataset, and compares it to other important models in the literature. Such a comparison is difficult, as different models are trained using different datasets, and have different restrictions for which samples can be processed. MHCAttnNet,^7^ while restricted to monoallelic DR samples, is available for retraining, and thus we can accurately measure its performance on our test dataset, without the worry of training data leakage. We observe an AP of 28.27%, compared to the 83.71% that our baseline graph-pMHC performs on the same monoallelic DR samples. On the other hand, NetMHCIIpan-4.0,^6^ while not available for re-training, is a pan-allelic, pan-allotypic model trained with a relatively up to date dataset, our baseline graph-pMHC significantly outperforms NetMHCIIpan-4.0 82.34% to 59.50%. MixMHCIIPred-1.2,^21^ also not available for re-training and limited to the monoallelic alleles that it is trained on, performs similarly (62.24%). Notably, our test data could potentially feature data points present in the training data of some of these methods but not in that of Graph-MHC. Despite this, the observed gain in performance is appreciable.

**Figure 4.**
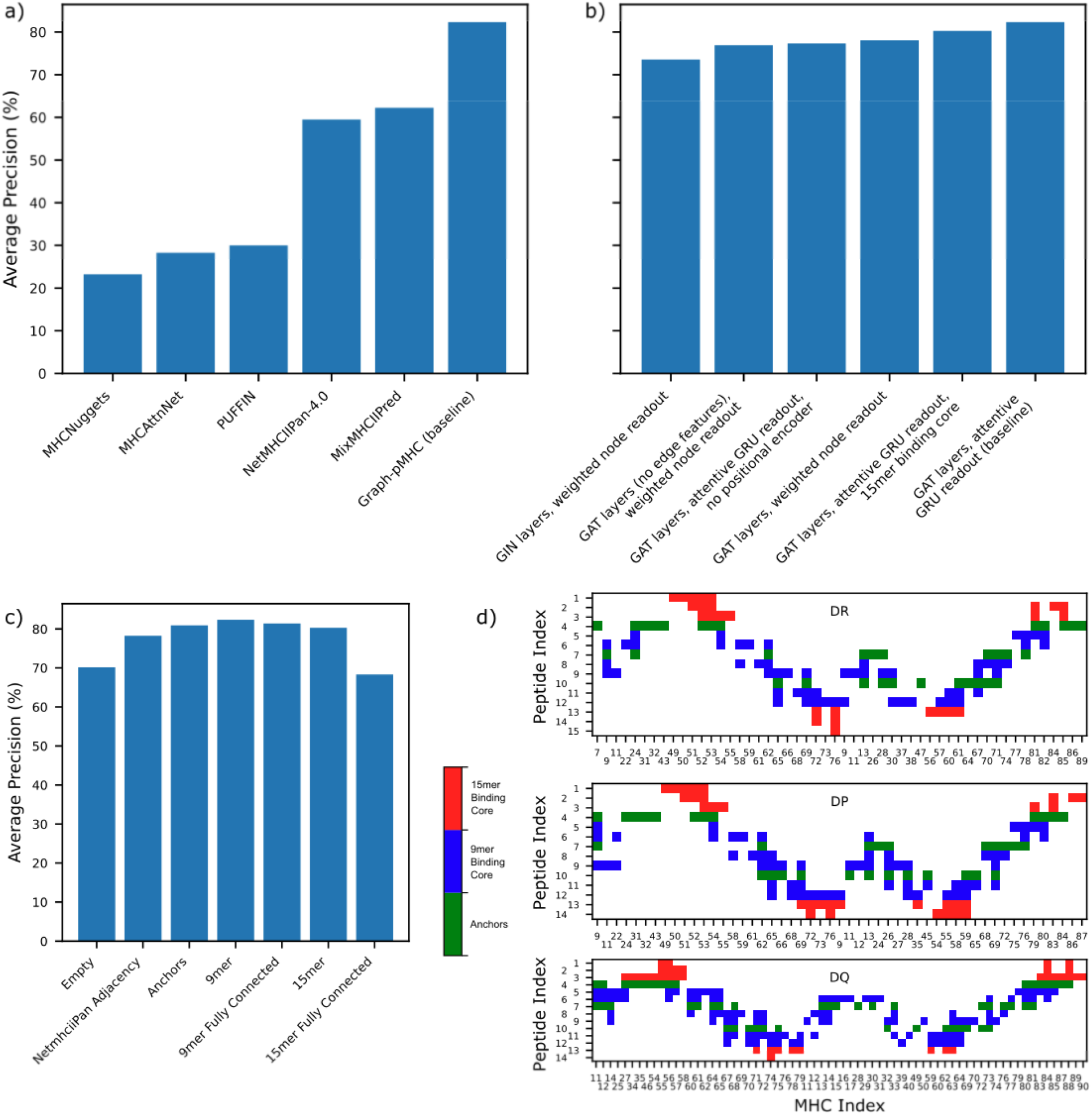
pMHCII presentation performance. a) Comparison of Graph-pMHC test set performance with other models found in the literature. All models are only evaluated on subsets of our test dataset for which inference can be performed. b) Impact of alternative model choices on test set performance. c) Barchart of test set performance with the various contact maps used to observe the impact of inductive bias on test set performance. d) Diagram of the various contact maps used.

In order to identify which features of graph-pMHC are the most important, we perform an ablation study, depicted in Figure 4 b). We find that choice of message passing layers is the most important factor in model performance, with graph attention layers (GAT)^22^ performing the best. Edge features are next most important; excluding it results in an AP drop of 5.33%. Positional encoding is similarly important, leading to a 5.04% AP drop. The choice of graph readout also contributes 4.28% over weighted node readout). The combination of attentive GRU GAT layers and attentive GRU graph readout makes the baseline graph-pMHC model similar to the AttentiveFP^23^ model developed for small molecule drug discovery. A key difference is the addition of positional encoding (and thus the hybrid graph-sequence strategy developed here).

We found that a 9mer binding core leads to a 2.06% AP decline over a 15mer binding core, despite the information lost. We further examine the impact of pMHCII adjacency matrix on performance in Figure 4 c). As expected, an empty adjacency matrix (with only intramolecular and flank edges included) dramatically reduces performance (12.17%), more than any ablation considered in Figure 4 b). We find that while anchor residues contribute most of the information in peptide motifs (for example Figure 5 a-c), they are not alone sufficient as including peptide-MHCII adjacencies for residues between anchors increases AP 1.41%. Still, we found that minimizing adjacencies was important: a fully connected binding core-MHCII adjacency matrix leads to a 0.96% AP drop.

**Figure 5.**
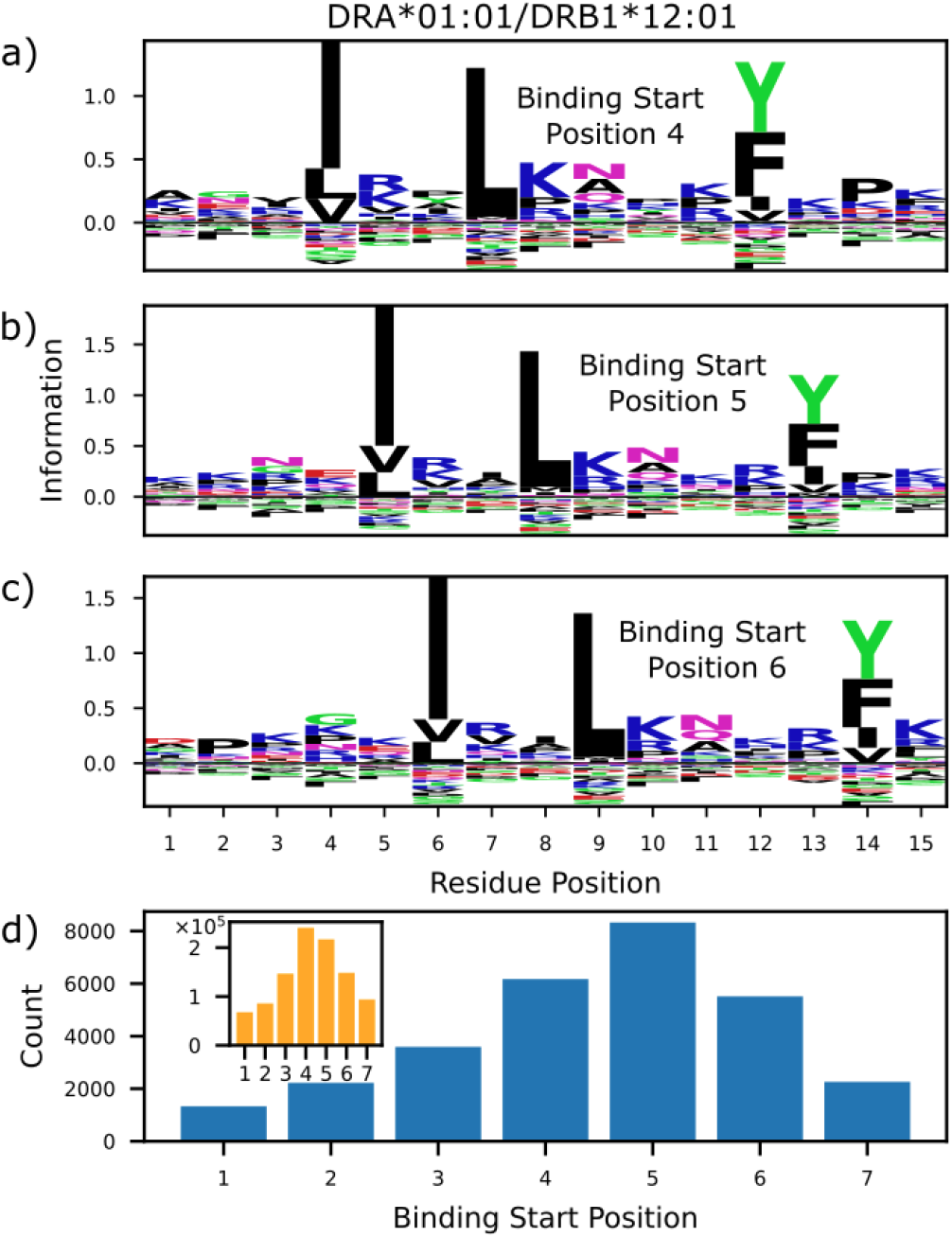
the importance of processing signals on graph-pMHC predictions. a-c) Peptide motifs of very likely presenters determined by graph-pMHC for DRA*01:01/DRB1*12:01. KL divergence of residue type with respect to observed residue frequency in humans for various binding core starting positions is depicted. b) Enrichment of likely binders for different starting positions, vs (inset) the overall number of binding core starting positions.

Graph-pMHC can also be used to understand the underlying biology in the pMHCII presentation pathway. To this end, we investigate peptide processing signals uncovered by the model using the allotype DRA*01:01/DRB1*12:01. Figure 5 a-c) depict motifs derived from the most likely 0.1% presenters from random 15mer peptides. One may observe the conservation of the binding core motif regardless of binding core starting position. Interestingly though, one can observe a consistent lysine and proline enrichments in peptide positions 15 and 14, respectively. The conservation of this enrichment, regardless of binding core starting position was measured experimentally in Barra et al,^24^ and suggests that these are signals to processing enzymes to cut the peptide at these positions. The model also predicts a strong enrichment of peptides whose peptide flanking regions hang off the MHC binding pocket preferentially in the N direction, as depicted in Figure 5 d). One can observe a preference in later binding core starting positions, until positions 6 and 7, where incompatible enrichment of anchors and processing signals leads to a precipitous drop.

### Application to Antibody Immunogenicity

Deimmunizing antibody drugs is a key application of pMHCII models, and so we have curated a dataset of 107 antibodies (AB) with the observed antidrug-antibody (ADA) response observed in published clinical trials. To evaluate the ADA risk of antibodies, we must first develop a method of summarizing the ADA risk of an AB from the peptide constituents. An additional difficulty in the evaluation is that t-cells capable of recognizing self cells will be negatively selected to prevent an autoimmune response, so even if a peptide is presented, it may not lead to an immune response.^1^ Thus, we must also remove self-peptides from the evaluation. Figure 6 a) depicts our approach for creating an AB ADA risk. First, presentation scores and binding cores are obtained for all length 12-19 peptides which can be derived. Then peptides whose binding cores do not differ from the nearest germline sequence in IMGT are removed. Additionally, we remove peptides whose binding cores appear 3 times in OAS.^25^ Next, peptides with a presentation logit score under 0 are removed (or likely binders for NetMHCIIpan-4.0), and the total number of unique binding cores obtained. This is repeated for 8 common DR alleles which were chosen to span most of the DR supertype families,^26^ and the total number of unique binding cores is used to represent the AB ADA risk.

**Figure 6.**
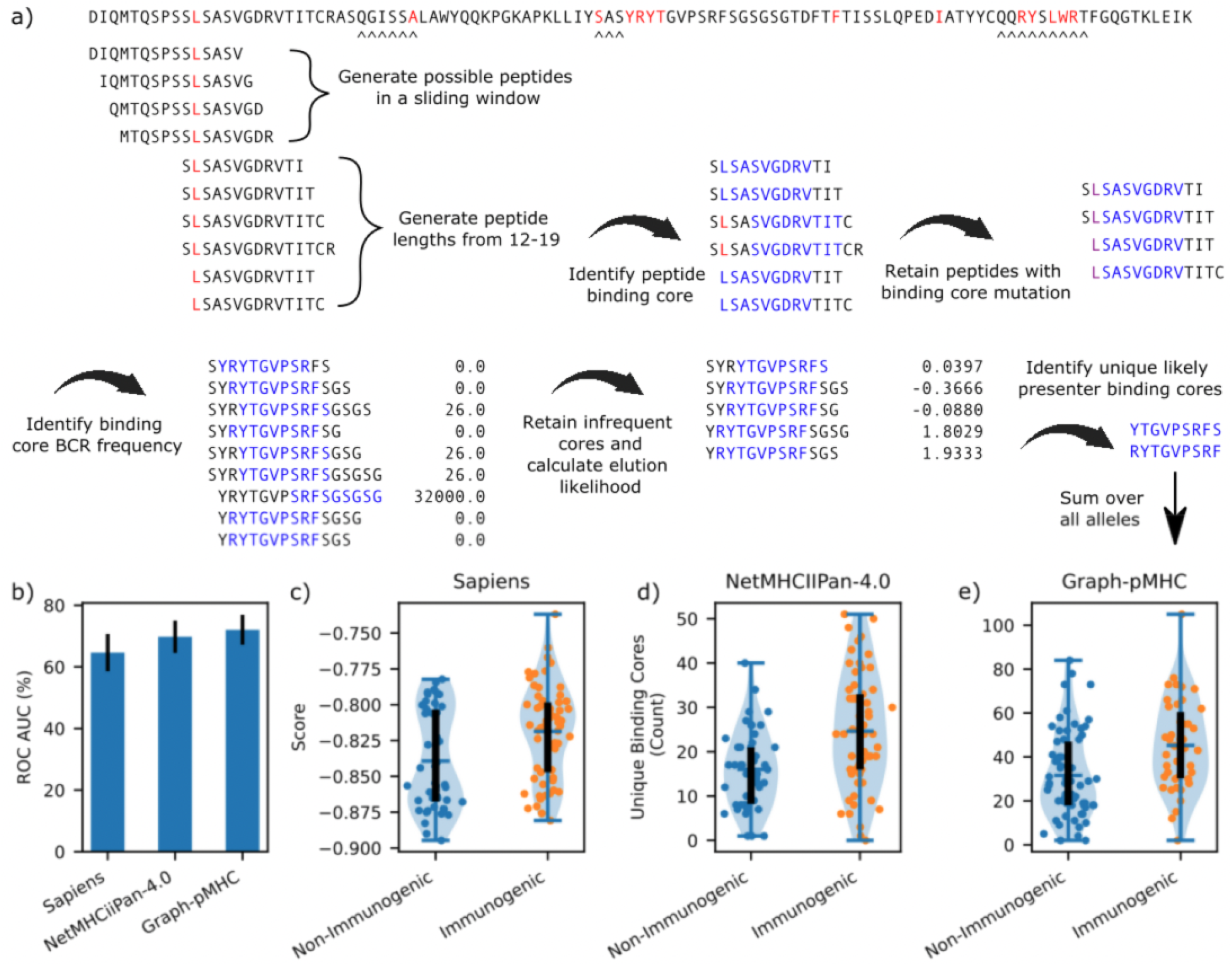
application of pMHCII to AB ADA risk assessment. a) Schematic depicting how ADA risk of an AB is calculated. All derivable peptides are obtained, and then various filters are applied to eliminate human-like peptides, and then the number of unique binding cores found in presentable peptides is counted. b) Barchart of the bootstrapped (500) ROC AUC for the models tested. c-e) Violin plot of AB ADA risk for the ABs in the dataset. The blue bar represents the median value, the black line runs from the 1st to 3rd quartile, blue and orange dots represent a non-immunogenic, and immunogenic AB, respectively. Sapiens (c) measures immunogenicity with an average score, higher is more immunogenic, while NetMHCIIPan-4.0 (d) and Graph-pMHC (e) use the strategy depicted in (a), where more unique binding cores is more immunogenic.

Figure 6 b) depicts the ROC AUC achieved by graph-pMHC and NetMHCIIpan-4.0. In this task, only a small improvement (2.25%) is observed using graph-pMHC over NetMHCIIpan-4.0. We attribute this to the fact that the task is merely to identify binding cores, and not rank peptides, as in presentation ranking considered above. Regardless, both models achieve clinically useful separation between the immunogenic and non-immunogenic ABs, as observed for NetMHCIIPan-4.0 and graph-pMHC in Figure 6 d), and e), respectively. We compare the strategy we developed here using pMHCII presentation prediction models to Sapiens, a transformer-based deep learning model trained to similarity of ABs with human derived BCRs. We observe a larger improvement in performance (7.41%) using pMHCII presentation models compared to Sapiens, which gives likely immunogenic predictions for a subset of non-immunogenic antibodies.

## Discussion

At its core, the task of predicting which peptides will be presented by MHC class ii is a biophysical one which is determined by the interactions of amino acid residues between the peptide and MHC allotype. Graph neural networks (GNN) are uniquely capable of capturing the interaction behavior by modeling adjacent residues in the peptide-MHCII complex (pMHCII) using graph edges, thus only mixing node information of adjacent residues. Due to this, we believe that GNNs should have a superior inductive bias for the pMHCII presentation than the popular sequence-based and multilayer perceptron-based methods that currently dominate the field.

Our analysis of pMHCII structures suggests a few constraints for a GNN approach which are born out in the ablation study conducted in Figure 4 b. First, a model must be able to perform an alignment of the peptide binding core relative to MHCII, to address this challenge we introduce our graph enumeration strategy. Second, while adjacencies are strongly conserved for different pMHCII, significant variations occur. This suggests a graph attention based approach (shown to be the most performant message passing framework in our tests), where the model can silence deleterious interactions. Third, the limited diversity in pMHC graphs yields a system somewhere between traditional sequence based machine learning and graph based machine learning. Due to this, positional encoding is applicable and significantly boosts performance, partly by helping inform the model about shifting binding cores. Finally, as binding is driven primarily by anchor residues, a GNN approach must have a sophisticated way of altering the representation of one anchor based on another, despite the fact that the anchors have no connecting edges. Indeed, we find an attentive gated recurrent unit (GRU) graph readout adds significant value compared to simple weighted node readouts.

The primary motivation of a GNN approach is the inductive bias that they bring. Unlike other methods, GNNs enable us to directly inform the model about which residues should exchange information with one another. In Figure 4 a), we benchmark the performance of popular models in the literature with a broad array of inductive biases. Interestingly, we find that the convolutional neural network (CNN) approach, PUFFIN, outperformed the recurrent neural network (RNN) approaches of MHCAttnNet and MHCNuggets. Yet, all three approaches significantly underperform graph-pMHC, demonstrating the utility of the GNNs structure-informed inductive bias for the task.

We further probe the impact of injected prior knowledge (and thus inductive bias) on performance by analyzing various adjacency matrix choices in Figure 4 c). The single most impactful element of prior knowledge was the 9mer binding core (and graph enumeration alignment). Yet we found that, even relying on graph attention to silence unnecessary adjacencies, overspecification of interactions begins to hamper performance. For this reason, the 15mer binding core, which was identified by alphafold2-multimer (AF2), underperforms the 9mer binding core, indicating that the residues on the outside flank of the anchors are not informative of peptide binding. Underspecification too was a significant issue, with an empty pMHCII adjacency leading to much poorer performance, and only specifying the anchor residues leading to a performance decline. We see then that the AF2 derived adjacency matrix strikes the ideal balance for inductive bias in this setting.

While the all-to-all residue inductive bias provided by multilayer perceptrons (MLPs) has been profitable for NetMHCIIpan-4.0, in our analysis of peptide processing in Figure 5 a), we show that dropping residues to satisfy the fixed input size requirement of the MLP may lead to dropping processing signals. To our knowledge no previous model has reported lysine/proline enrichment outside of the binding core, and it indicates the sensitivity of the graph-pMHC approach. This is particularly relevant for very long peptides where these enriched residues may be distant from the 15mer input used for NetMHCIIpan-4.0 as these residues will not be included as input.

Finally, we provide a new dataset for evaluating antibody immunogenicity, which is mediated by CD4 t cells. Previous datasets were almost completely composed of mouse derived antibodies, limiting their usefulness in modern antibody engineering. By obtaining more human derived antibodies, we enable the evaluation of methods for obtaining antibody immunogenicity. Indeed, the previous antibody immunogenicity dataset used by Sapiens led to no statistically significant separation of immunogenic and non-immunogenic human antibodies by their model, where good performance of their model is observed on our dataset.^13^ Further, we develop a strategy for using pMHCII presentation models to create an AB immunogenicity risk score, which outperforms Sapiens human-ness based score. As complementary sources of information, antibody engineers can use both graph-pMHC and Sapiens to guide the humanization of antibodies. This application of graph-pMHC has the potential to increase the success rate of clinical trials, which often fail due to immunogenicity.

## Methods

### Contact map generation with Alphafold2-mutlimer (AF2)

The 5 most likely to present peptides for each allele (as chosen by a graph-pMHC model with NetMHCIIpan adjacency), along with the top peptide for each peptide length shown in Figure 1 d) are used to derive adjacency matrices. These sequences, along with the MHC sequences (taken from the Immunogenetics information system (IMGT)) are used to obtain predictions from AF2 on the default settings. Peptides which are predicted not to lay flat in the binding groove, or run N-C (opposite typical peptides) in the binding groove are filtered out. An adjacency matrix is constructed by identifying contacts within 4 Å between any atom on any peptide residue with any atom on any MHC residue. Peptides are aligned by the residue which is in contact with the 8th MHC residue (or 10th MHC residue if no contact is observed with the 8th residue). The canonical gene adjacency matrix is then defined using contacts which occur at least 20 times for DR, and 10 times for DP and DQ.

### Graph Generation

The AF2 contact map is used to define bidirectional intermolecular edges between peptide and MHC, adjacent residues in each sequence are given bidirectional intramolecular edges (MHC residues not in contact with peptide are not included in the graph to save computation, and nearest neighbors in the psuedosequence are given intramolecular edges). Bidirectional flank edges are made between the n-most residue in the protein n-flank and the c-most residue in the peptide c-flank. The c-most residue of the protein c-flank and the n-most residue in the peptide n-flank are also linked in this way. Intermolecular, intramolecular, and flank edges are defined as one-hot encoded vectors. To amortize the graph generation process, all possible graphs are generated once, and a lookup table is used to assign flank-peptide-flank-MHC sequences to the appropriate graphs, which is fully defined by the lengths of the peptide, flanks, MHC gene, and binding core starting position.

### pMHCII presentation model implementation

Graph-pMHC is depicted diagramatically in Figure 2 c, and algorithmically below. Input into the model is a batch of the various sequences, n-flank, peptide, c-flank, and up to 12 allotypes. First, n-flank, peptide, c-flank, allele pairs are generated for each allele. Node features are generated for each residue (empty flanks are given a special token) in these pairs via nn.embedding (dim 64), and positional embedding is applied by adding a unique token per position which is embedded via nn.embedder (dim 64). Graphs and edge features are looked up for each pair for every possible binding core starting position, and these are used in GAT layers (dim 128, n_layers 2, dropout 0.1) to update the node features. These node features are sent through an attentive GRU, described in AttentiveFP^23^ (dim 128, time_steps 2, dropout 0.2), to generate graph features. These graph features are sent through a linear layer to one output node which captures the presentation likelihood. The best binding core starting position and allele for each input peptide is found, and loss is calculated using that combination. Graph-pMHC is implemented in pytorch and deep graph library, and uses the fast.ai library for constructing its training loop. As with previous work, we use negative set switching between epochs.^27^

### Train/Test split development

To minimize 9mer overlap, we began with a set of all protein sequences within the proteome (Ensembl v90). For each Ensembl gene, we generated a set of unique 9mers from all of their associated Ensembl proteins. From these sets of unique 9mers we generated a table of the most common 9mers across Ensembl genes. We sought to place the most common kmers from the proteome within the training set, while retaining more unique kmers for the test set. To quantify overlap, we measured the number of overlapping 9mers between train and test relative to the number of unique 9mers.

To minimize skew of protein function, process, and localization between train and test we leveraged Gene Ontology (GO) information. GO domains (molecular function, biological process, and cellular compartment) and terms were mapped to Ensemble genes and limited to GO terms containing at least 10 member genes (given our desired 9:1 split, smaller terms would be impossible to divide evenly). To measure evenness of GO term distribution, we measured the R^2^ of the proportion of each GO term within train and test.

To integrate and balance these two objectives (minimized kmer overlap and relatively even GO term distribution), we implemented an iterative process to separate genes into train and test. At a high level, we iterated through each GO term, separating genes associated with that term into train and test based on whether they contained globally frequent proteome kmers. This process continued across GO terms until all genes had been placed. More specifically, beginning with the smallest GO term in the molecular function domain, we took all genes associated with that term. Then beginning with the most abundant kmer in the global kmer table, added all gene’s containing that kmer to the list of training genes. We continued this process down the global kmer list until >= 90% of genes associated with the GO term were found in the list of training genes. We repeated this process each GO term in the molecular function domain, accounting for genes that had been assigned in previous iterations as well. We then repeated this process for GO terms in the biological process and cellular component domains and finally for the genes that had no GO term annotation.

### Antibody immunogenicity risk assessment

Our strategy for antibody (AB) immunogenicity risk assessment is depicted diagramatically in Figure 6 a). All peptides with length 12-19 are generated in a sliding window across each AB (with flanks), these peptides are paired with DRB1*01:01, DRB1*03:01, DRB1*04:01, DRB1*07:01, DRB1*08:01, DRB1*11:01, DRB1*13:01, DRB1*15:01. Graph-Pmhc is used to obtain elution likelihood scores and binding core starting predictions for each peptide-allele pair, and pairs with score less than 0 are filtered out (for NetMHCII-pan, peptides rated less than weak binders are filtered out). The binding core frequency dataset is created from OAS^25^. Human PBMC, no disease, no vaccine BCRs are extracted from OAS and limited to just productive vdj complete BCRs. This leaves 39,244,222 heavy, 8,077,461 lambda, and 109,138 kappa chains. All 9mers are extracted and their frequencies are recorded. Graph-pMHC identified binding cores from the AB dataset are filtered out if they appear 3 times in this list of BCR 9mers. Additionally, binding cores which do not contain a residue which is different from the chain’s nearest neighbor is also filtered out, as determined using the ABnumber library. The number of unique binding cores per allele is summed up, and this is used as the immunogenicity risk.

