## Supplemental Information for "Graph-pMHC: Graph Neural Network Approach to MHC Class II Peptide Presentation and Antibody Immunogenicity"

**Supplementary Information**

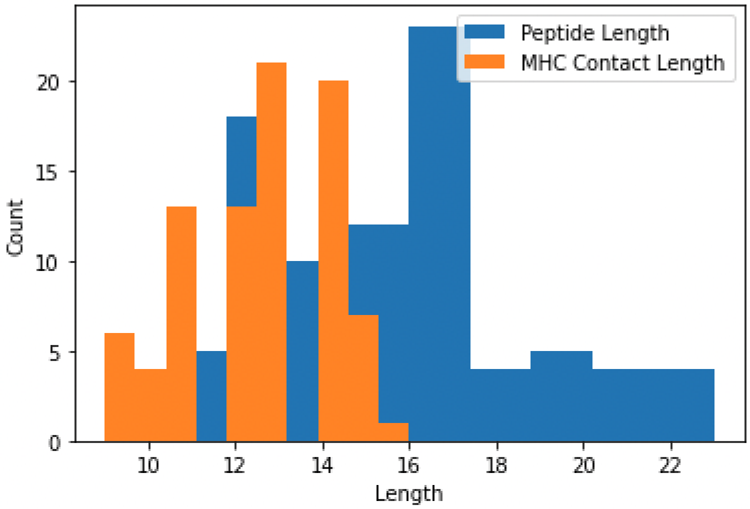


Supplemental Figure 1 depicts the distribution of peptide lengths used to obtain AF2 structures (blue) and the resulting distribution of lengths in contact with MHC2 (orange).

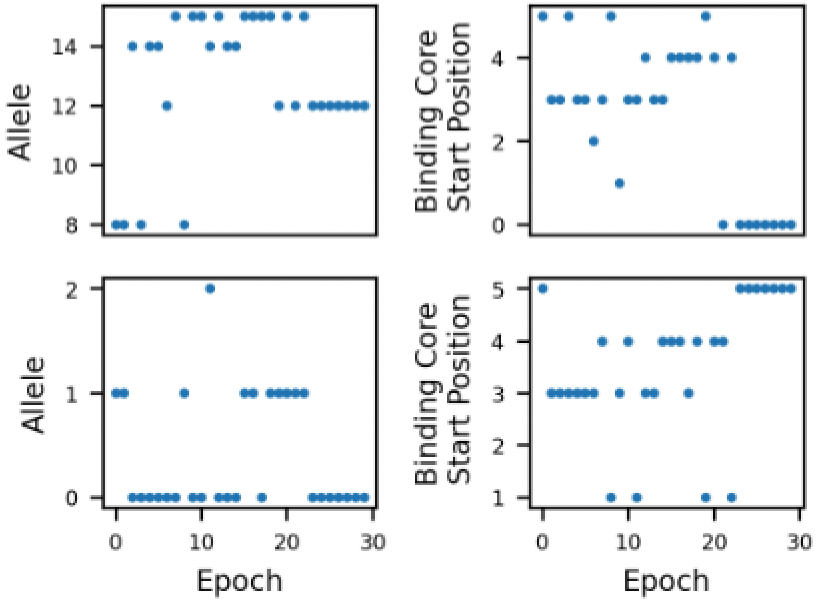


Supplemental Figure 2 depicts the convergence of allele choice and binding core starting position for two particularly difficult peptide:genotype pairs (left: DSSVPRNKNPFQEA:DPA1*01:03___DPB1*04:01_DQA1*01:01_DQA1*05:05_DQB1*03:01_DQB1*05:03_DRA*01:01_DRB1*11:01_DRB1*14:54_DRB3*02:02 right: LSALEEYTKKLNTQ:DRA*01:01_DRB1*13:01_DRB1*14:01_DRB3*01:01_DRB3*02:02).
